# SparseRNAFolD: Sparse RNA pseudoknot-free Folding including Dangles

**DOI:** 10.1101/2023.06.05.543808

**Authors:** Mateo Gray, Sebastian Will, Hosna Jabbari

## Abstract

**Motivation:** Computational RNA secondary structure prediction by free energy minimization is indispensable for analyzing structural RNAs and their interactions. These methods find the structure with the minimum free energy (MFE) among exponentially many possible structures and have a restrictive time and space complexity (*O*(*n*^3^) time and *O*(*n*^2^) space for pseudoknot-free structures) for longer RNA sequences. Furthermore, accurate free energy calculations including dangles contributions can be difficult and costly to implement, particularly when optimizing for time and space requirements.

**Results:** Here we introduce a fast and efficient sparsified MFE pseudoknot-free structure prediction algorithm, SparseRNAFolD, that utilizes an accurate energy model that accounts for dangles contributions. While sparsification technique was previously employed to improve time and space complexity of a pseudoknot-free structure prediction method with a realistic energy model, SparseMFEFold, it was not extended to include dangles contributions due to complexity of computation. This may be at the cost of prediction accuracy. In this work, we compare three different sparsified implementations for dangles contributions and provide pros and cons of each method. As well, we compare our algorithm to LinearFold, a linear time and space algorithm, where we find in practice, SparseRNAFolD has lower memory consumption across all lengths of sequence and a faster time for lengths up to 1000 bases.

**Conclusion:** Our SparseRNAFolD algorithm is an MFE-based algorithm that guarantees optimality of result and employs the most general energy model including dangles contributions. We provide basis for applying dangles to sparsified recursion in a pseudoknot-free model which has the ability to be extended to pseudoknots.

**Availability:** SparseRNAFolD’s algorithm and detailed results are available at https://github.com/mateog4712/SparseRNAFolD.

## 1 Introduction

Non-coding RNAs play crucial roles in the cell such as in transcription [3], translation [3, 17], splicing [24, 33], catalysis [3, 39] and regulating gene expression [3, 12, 19, 24]. Since RNA’s function heavily relies on its molecular structure, facilitated by hydrogen bonding both within and between molecules, predicting and comprehending the structure of RNA is a dynamic area of research. It is reasonable to assume (without further knowledge) that RNA forms the structure with the lowest free energy [20, 25]. This is the motivation for algorithms that aim to predict RNA minimum free energy (MFE) structure from a pool of exponentially many structure it can form. Such methods employ a set of energy parameters for various loop types, called an energy model; to find the free energy of a structure, they add up the energy of its loops. While prediction accuracy of these methods depends on the quality of their energy models, these methods are applicable to novel RNAs with unknown family or function and for prediction of structure of interacting molecules. The large time and space complexity of MFE-based methods (*O*(*n*^3^) time and *O*(*n*^2^) space where *n* is the length of the RNA), however, restricted their applications to small RNAs. Sparsification technique was utilized recently to existing MFE-based algorithms to reduce their time and/or space complexity [36, 16, 14, 35, 30, 23, 2, 4] by removing redundant cases in the complexity-limiting steps of the dynamic programming algorithms. While majority of these methods focused on simple energy models, some expanded sparsification technique to more realistic energy models [36, 16, 14]. To the best of our knowledge, no existing method has yet incorporated dangles energy contributions into a sparsified prediction algorithm. Dangle energies refer to the free energy contributions of unpaired nucleotides that occur at the end of a stem-loop structure.

We show in Figure 1 the location of dangles on a pseudoknot-free structure (see Figure 1a) and a pseudoknotted structure (see Figure 1b). The complexity of dangles in a pseudoknot further occurs as dangles have to be tracked for both bands within the pseudoknot.

**Figure 1.**
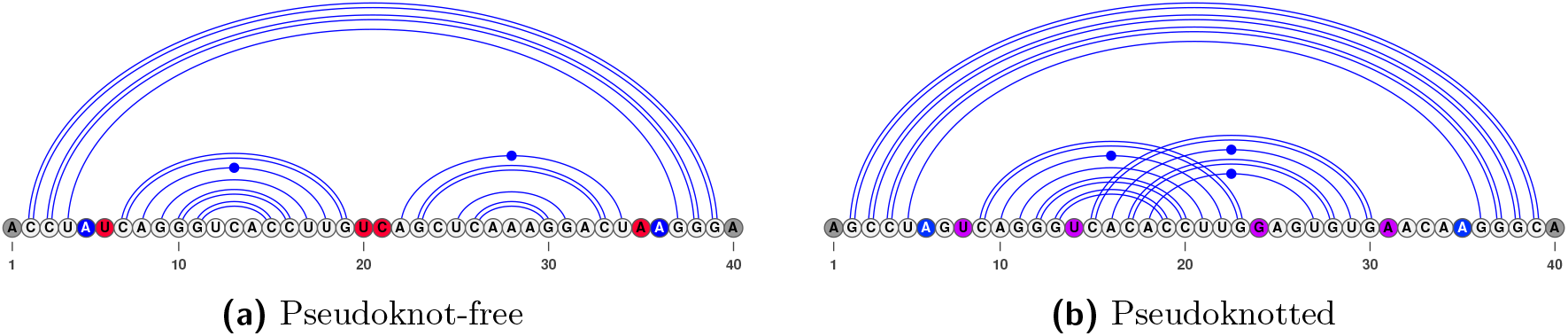
An RNA structure is shown with dangles highlighted. **(a)** In red, we have the dangles on the bands in the multi-loop. In blue, we have the dangle on the closing bases of the multi-loop. In gray, we have dangles on the outer end of the RNA. **(b)** We include purple to show dangles occurring in a pseudoknot. Dangles in pseudoknots can be handled differently depending on the program.

Neglecting dangle energies in the prediction of RNA structure stability can lead to inaccuracies. For instance, a stem-loop structure that includes an unpaired nucleotide at the end may appear less stable than its actual stability if the dangle energy contribution is ignored. Conversely, a stem-loop structure with an unpaired nucleotide that interacts positively with another one may appear more stable than its actual stability if the dangle energy contribution is not taken into account.

Dangles, in some form, are implemented in the majority of MFE pseudoknot-free secondary structure prediction algorithms [9, 8]. RNAFold [9, 11, 22, 6, 18, 45, 7] is an *O*(*n*^3^) time and *O*(*n*^2^) space algorithm which implements the dangle 0 (“no dangle”), dangle 2 (“always dangle”), and dangle 1 (“exclusive dangle”) model (defined in Section 2.1). It also utilizes a dangle model which implements coaxial stacking – a type of stacking which gives a bonus to stacks in the vicinity of each other. LinearFold [8], a sparsified *O*(*n*) space heuristic algorithm has implemented the “no dangle” and “always dangle” model but has not implemented an “exclusive dangle” model. Fold from the RNAstructure library [28] is an *O*(*n*^3^) time and *O*(*n*^2^) space algorithm which implements an “exclusive dangle” model with coaxial stacking. MFold [44, 34, 43] is an *O*(*n*^3^) time and *O*(*n*^2^) space algorithm which has implemented an “exclusive dangle” model with coaxial stacking.

Handling dangles in pseudoknot prediction algorithms are less developed. Pknots [29], an *O*(*n*^6^) time and *O*(*n*^4^) space pseudoknot prediction algorithm has implemented an “exclusive dangle” model which also includes coaxial stacking. Within Pknots, a set of parameters are defined for non-pseudoknot and pseudoknot dangles. The pseudoknot parameters are estimated and rely on an estimated weighting parameter. Hotknots [27], a heuristic algorithm, uses the DP09 parameters which includes pseudoknotted parameters from Dirks and Pierce [5] and tuned by Andronescu et al. [27]; however, the energies for the pseudoknotted dangles are the same as pseudoknot-free dangles and there is no weighting parameter.

## Contributions

In [37, 36], we already discussed the sparsification of RNA secondary structure prediction by minimizing the energy in the Turner energy model. However, in this former work we did not yet consider the energy contributions due to the interactions of base pairs at helix ends with dangling bases (i.e. ‘dangling ends’). Here, we identify the correct handling of dangling end energies in the context of sparsification as a non-trivial problem, characterize the issues, and present solutions.

For this purpose, we first state precisely how dangle energies are handled by energy minimization algorithms; to the best of our knowledge, this is elaborated here for the first time. Consequently, we devise novel MFE prediction algorithms that include dangling energy contributions *and* use sparsification techniques to significantly improve the time as well as space complexity of MFE prediction.

Like the algorithm of [36], our efficient SparseRNAFolD algorithm keeps the additional information to a minimum using garbage collection. In total, we study three different possible implementations and compare their properties which make them suitable for different application scenarios. Finally, while we study the case of non-crossing structure prediction, we discuss extensions to the more complex cases of pseudoknot and RNA–RNA interaction prediction (such extensions being the main motivation for this work in the first place).

## 2 Preliminaries: sparsification without dangling ends

We restate the preliminaries and main results from our former work on sparsification of free energy minimization without dangling ends [36].

We represent an ***RNA sequence*** of length *n* as a sequence *S* = *S*_1_, …, *S*_*n*_ over the alphabet {*A, C, G, U*} ; *S*_*i,j*_ denotes the ***subsequence*** *S*_*i*_, …, *S*_*j*_. A ***base pair*** of *S* is an ordered pair *i*.*j* with 1≤ *i* < *j*≤ *n*, such that *i*th and *j*th bases of *S* are complementary (i.e. {*S*_*i*_, *S*_*j*_} is one of {*A, U*}, {*C, G*}, or {*G, U*}). A ***secondary structure*** *R* for *S* is a set of base pairs with at most one pair per base (i.e. for all *i*.*j, i’ .j’* ∈*R*: {*i, j*} ∩ {*i ‘, j’*} =∅). Base pairs of secondary structure *R* partition the unpaired bases of sequence *S* into ***loops*** [26] (i.e., hairpin loop, interior loop and multiloop). Hairpin loops have a minimum length of *m*; consequently, *j*− *i* > *m* for all base pairs *i*.*j* of *R*. Two base pairs *i’*.*j’* and *i .j* cross each other iff *i*’ < *i*’ < *j*< *j*’ or *i*’ < *i* < *j*’ < *j*. A secondary structure *R* is ***pseudoknot-free*** if it does not contain ***crossing base pairs***.

The unsparsified, original algorithm for energy minimization over pseudoknot-free secondary structures was stated by Zuker and Stiegler [42]. It is a dynamic programming algorithm that, given an RNA sequence *S* of length *n*, recursively calculates the minimum free energies (MFEs) for subsequences *S*_*i,j*_ as *W* (*i, j*) (stored in a dynamic programming matrix).

### SparseRNAFolD

Finally, *W* (1, *n*) is the optimal free energy. We state this algorithm in a sparsification-friendly form following [36]. As usual, the algorithm is described by a set of recursion equations (for a minimum hairpin loop size *m* and a maximum interior loop size *M*). For 1 ≤ *i* < *j* ≤ *n, i* < *j* − *m*:

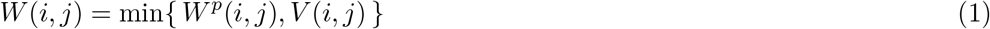

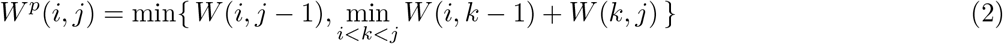

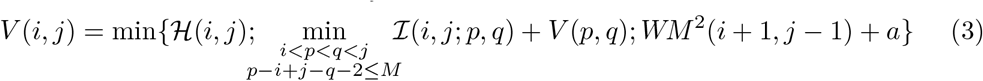

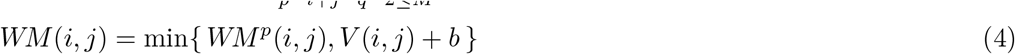

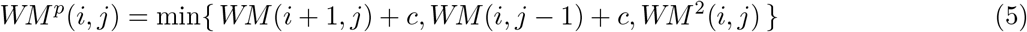

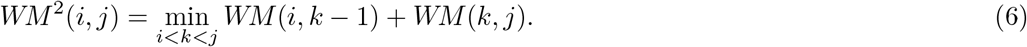

Here, *a, b, c* are multi-loop initialization penalty, branch penalty and unpaired penalty in a multi-loop, respectively. The initialization cases are *W* (*i, i*) = 0; *V* (*i, j*) = *WM* (*i, j*) = ∞ for all *j* − *i* ≤ *m* and *WM* ^2^ = ∞ for all *j* − *i* ≤ 2*m* + 3.

In these recursions, all function values (like *W* (*i, j*) or *W*^*p*^(*i, j*)) denote minimum free energies over certain classes of structures of subsequences *S*_*i,j*_. The classical Zuker/Stiegler matrices *W, V* and *WM* are defined as: *W* yields the MFEs over general structures; *V*, over closed structures, which contain the base pair *i*.*j*; *WM*, over structures that are part of a multi-loop and contain at least one base pair.

Since sparsification is based on the idea that certain optimal structures can be decomposed into two optimal parts, while others (namely closed structures) are non-decomposable, we single out the partitioning cases and introduce additional function symbols *W*^*p*^, *WM*^*p*^, and *WM* ^2^.

### Sparsification without dangling ends

This allows to cleanly explain the *key idea of sparsification* and consequently formalize it: to minimize over the energies of general structures in *W* (*i, j*)—note that there is another minimization inside of multi-loops which is handled analogously—the algorithm considers all closed structures *V* (*i, j*) and all others *W*^*p*^(*i, j*). Optimal structures in the latter class can be decomposed into two optimal structures of some prefix *S*_*i,k−*1_ and suffix *S*_*k,j*_ of the subsequence. Classically, the minimum is therefore obtained by minimizing over all ways to split the subsequence. Sparsification saves time and space since it is sufficient to consider only the splits where the optimum of the suffix *S*_*k,j*_ is not further decomposable (formally, where *W* (*k, j*) < *W*^*p*^(*k, j*)). Briefly (for more detail see [37] or [35]), this is sufficient since otherwise there is a *k* to optimally split the suffix further into *S*_*k,k*_’ *−*1 and *S*_*k*_ ’,*j*. The split of *S*_*i,j*_ at *k* cannot be better than the split at *k* and therefore does not have to be considered in the minimization; thus, it can be restricted to a set of *candidates*. This argues by the ***triangle inequality for*** *W* (which directly follows from the definition of *W* as minimum):

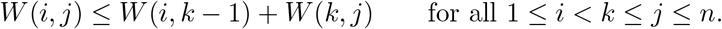

Consequently, sparsification improves the computation *W*^*p*^, *WM*^*p*^ and *WM* ^2^. The corresponding sparsified version are

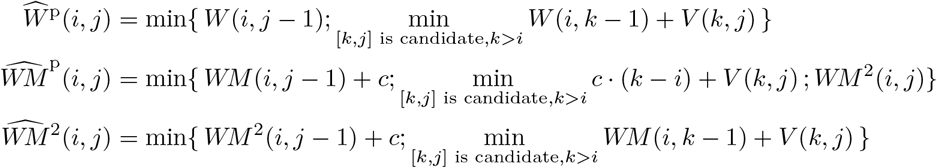

where candidates [*k, j*] correspond to the not optimally decomposable subsequences *S*_*k,j*_ (in either situation: general structures or structures inside of multi-loops), i.e. [*i, j*] is a ***candidate*** iff 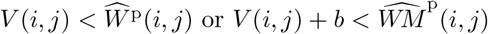

### Time and space complexity of sparsified energy minimization

Will and Jabbari showed that following the above algorithm *W* (1, *n*) can be calculated in *O*(*n*^2^ + *nZ*) time, where *Z* is the total number of *candidates*. While the MFE structure in the Zuker and Stiegler algorithm can be trivially reconstructed following a traceback procedure, this is not the case if sparsification is used for improving time *and* space as in the SparseMFEFold algorithm (and our novel algorithms). To improve the space complexity, sparsification avoids storing all entries of the energy matrix. The idea is to store the candidates and as little additional matrix entries as possible. A specific challenge is posed by the decomposition of interior loops (the single most significant major complication over base pair maximization, see [2]). For this reason, Will and Jabbari introduced *trace arrows* for cases, where the trace cannot be recomputed efficiently during the traceback procedure; they discussed several space optimization techniques, such as avoiding trace arrows by rewriting the MFE recursions, and removing trace arrows as soon as they become obsolete. Due to such techniques SparseMFEFold requires only linear space in addition to the space for candidates and trace arrows; its space complexity is best described as *O*(*n* + *T* + *Z*), where *T* is the maximum number of trace arrows.

### 2.1 Dangles

Recall that sparsification was discussed before (e.g. in SparseMFEFold) only for the simplest and least accurate variant of the Turner model, namely the one without dangling ends contributions. Before we improve this situation, let’s look in more detail at dangling ends and different common ways to handle them. Specifically, we discuss different *dangle models* “no dangle” (model 0), “exclusive dangle” (model 1), and “always dangle” (model 2) as implemented by RNAfold of the Vienna RNA package (and available via respective command line options -d0, -d1, and -d2).

Dangling end contributions occur only at the ends of stems (either in multiloops or externally) due to stacking interaction between the closing base pair of the stem and one or both immediately adjacent unpaired bases. In contrast, dangling end terms are not considered within (interior loops of) stems by the energy model.

We present modified DP recursions in order to reflect precisely, where and how dangling ends are taken into account. Therefore, in preparation, let’s replace *V* in the Equations (1) and (4) of the free energy minimization recursions of Section 2 by a new function *V* ^d^. The dangle models differ in the exact definition of *V* ^d^.

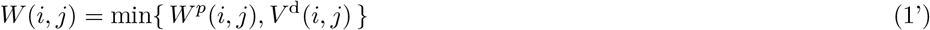

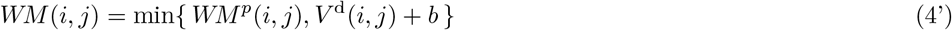

Note that in the energy model dangling ends can furthermore occur at the inner ends of helices that close a multi-loop. These dangles can be handled directly in the recurrence of *V* (*i, j*); specifically, in the subcase where *i*.*j* closes a multi-loop.

### No dangles

In the simplest model “no dangle”, dangling ends are ignored. We achieve this by defining

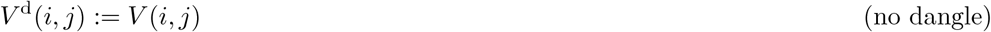

While easy to implement, it is clearly wrong to ignore dangling end contributions and this has a significant negative effect on the prediction accuracy compared to the other dangle models [31, 41, 40].

### Always dangle

A second relatively simple way is to apply a 53’ dangle energy at both ends of a stem (both 5’ and 3’ end), assuming that stem ends always dangle with their adjacent bases. As a strong simplification, in this model, one disregards whether the bases are paired and/or dangle with a different stem (either case would actually make them unavailable for dangling).

This dangle model allows the dangling ends to have a thermodynamic influence while keeping the model easy to implement as neither the conflicting adjacent nucleotides nor the energies of single dangle have to be tracked; it only requires knowledge of the bases on the 3’ and 5’ side of a base pair. Formally, we implement *V* ^d^ as

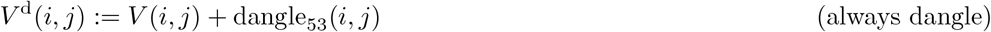

Moreover, we add the appropriate dangle contribution when closing a multi-loop in Eq. (3) in the last case of the *V* -recurrence of Eq. (3). The term *WM* ^2^(*i* + 1, *j* −1) + *a* is rewritten to

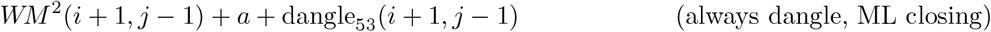

### Exclusive dangling

The most complex but general secondary structure dangle model “exclusive dangle” considers both single and double unpaired nucleotides adjacent to a stem. Furthermore, the model does not allow shared dangling ends i.e. no base can be used simultaneously in two dangles (in other words, adjacent unpaired bases dangle *exclusively* with a single stem end). As the restriction requires tracking of unpaired bases, *V* (*i, j*) places the possible unpaired bases at *i* and *j* and looks at the adjacent *V* energies. As this requires knowledge of energies adjacent to the current bases being looked at, this inherently causes a difficulty in sparsification.

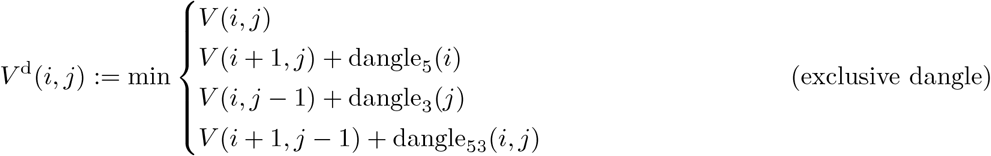

Moreover, we consider dangles at the closing of a multi-loop. In this model, the case *WM* ^2^(*i* + 1, *j*− 1) + *a* in the minimization of Eq. (3) is replaced by (the minimum of) four different cases:

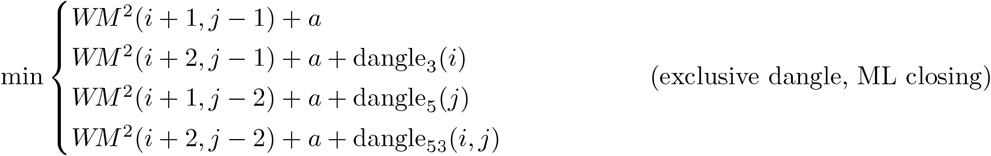

### 2.2 Space-efficient sparsification with exclusive dangles is non-trivial

We approach our main motivation for this work, which is to study and solve the issues of sparsification in the exclusive dangle model (dangle model 1). Let’s thus start by applying the idea of sparsification (Section 2) straightforwardly to the Recursion (2) (where *W* and *V* ^d^ are defined for exclusive dangles).

We quickly come up with the equation

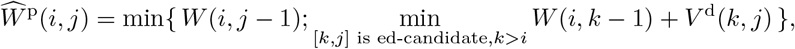

but would still have to define *ed-candidate* (exclusive dangle candidate) to make this work. We could define: [i,j] is an ***ed-candidate*** iff 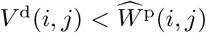, where the correctness of sparsification holds to a sparsification-typical triangle inequality argument (Section 2).

Expanding *V* ^d^ shows that this is not the only possible path to sparsifying the recursion. We could consider

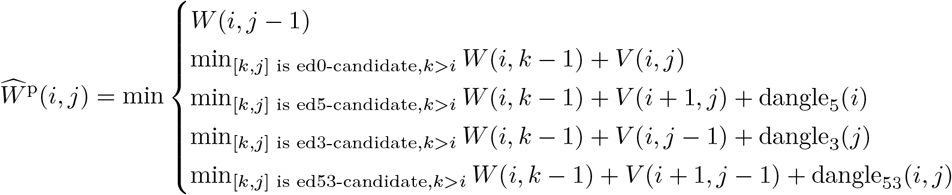

with different sets of candidates for all four cases. However, storing all these candidate sets (recall that there is even a second recursion that needs to be sparsified) is easily prone to compromise any space benefits due to sparsification in practice.

The transfer of the techniques from [36] brings even more problems, since due to such definitions, candidates [i,j] do not necessarily correspond to subsequences which closed optimal structure. Will and Jabbari strongly exploited this fact for their strong space savings.

Even considering our definition of an ed-candidate, we still run into the challenge to trace back to the corresponding base pair. With just the dangle energy, this poses issues as an ed-candidate can be one of four cases.

#### Lemma 1.

*In the exclusive dangle model, storing only the energy of each ed-candidate is not sufficient to correctly trace back from the candidate*.

**Proof**. Concretely, for the loop-based Turner 2004 energy model [21]) with exclusive dangles, consider the following RNA sequence *S* of length 12 with its MFE structure:

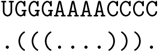

In the calculation of *W* (1, 12), the recurrences unfold to *W* (1, 12) = *W* (1, 1) + *V* ^d^(2, 12) = *W* (1, 1) + *V* (2, 11) + dangle_3_(12) =… = −2.9 kcal/mol, i.e. it is optimal to assume dangling of base pair (2, 11) to the right.

In a non time- and space-sparsified algorithm, recomputing *V* ^d^ from *V* adjacent energies would be trivial. However, due to space sparsification, the values of *V* are generally unavailable in the trace-back phase. In the constructed example, recomputation would require to know *V* (2, 12), *V* (2, 11), *V* (3, 12), and *V* (3, 11). Thus, under the assumption of the lemma, the optimal dangling cannot be efficiently recomputed for a candidate like [2,12]. ◂

In our preceding work SparseMFEFold [38], trace arrows were introduced to trace back to non-candidate values necessary to the structure within the interior loop case: *V* ^il-cand^(*i, j*). Trace arrows that point to candidates are not stored as they can be avoided by minimizing over candidates as seen in Equation 7.

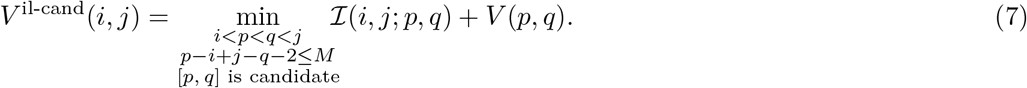

Consequently, finding the inner base pair of a loop through a candidate relies on the energy saved being *V* (*p, q*). However, as shown in Eq. (exclusive dangle, ML closing), the dangle energy could be *V* (*p, q*), *V* (*p*+1, *q*), *V* (*p, q* −1), or *V* (*p*+1, *q*− 1). Replacing the stored energy within a candidate with *V* ^d^ may conflict with the interior loop calculation. Recomputation of the *V* values required for *V*_*d*_ would negate the sparsification benefit. In summary, there is no easy or direct way to save the *V* energy required for the interior loop as well as the *V* ^d^ energy required for a multi-loop or external loop within the current candidate structure.

#### Lemma 2.

*The minimization over inner base pairs in the recursion of V cannot be restricted to candidates in the same way as in SparseMFEFold*.

**Proof**. Again consider the loop-based Turner 2004 energy model. There is a sequence *S* and

1≤ *i* < *j* ≤*n*, such that *V* ^d^(*p, q*) < *V* (*p, q*) but there is no way to trace back to *p* and *q* from *i* and *j*, namely consider the RNA sequence *S* of length 19 with its MFE structure:

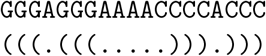

The optimal recursion case of *V* (3, 17) forms the interior loop closed by 3.17 with inner base pair 5.15, because *V* (5, 15) = −2.4 kcal/mol and V(3,17) = *I* (3, 17; 5, 15) + *V* (5, 15) = −1.5 kcal/mol.

The space optimization of SparseMFEFold removes trace arrows to candidates since the trace-back to candidates can be reconstructed based on candidate energies (compare Eq. (7). In the way of SparseMFEFold, we would not store a trace arrow pointing to 5.15 from [3, 17], since [5, 15] is a candidate. However, without a trace arrow, we would not reconstruct the correct trace. This happens, since the optimal structure in the subsequence 5. 15 GGGAAAACCCC would be (((….))). due to the 3’ dangle (*V* ^d^(5, 15) = −2.9 kcal/mol). Consequently, tracing back the optimal path from *V* ^d^(5, 15) wrongly introduces a base pair at 5.14. ◂

## 3 SparseRNAFolD

SparseRNAFolD combines the power of sparsification and a general energy model including dangle energies to achieve a fast and highly accurate RNA pseudoknot-free secondary structure prediction. To this end, we started from the sparsified dynamic programming recurrences of SparseMFEFold (which implements “no dangles” model), rewriting and revising them to accommodate various dangle energies.

### 3.1 “always dangle” model

Recall that “always dangle” model considers both 5’ and 3’ ends of a branch of a multi-loop or external loop for dangles contributions. Addition of this model is trivial with no change necessary to the recurrences of the SparseMFEFold. Note that as mentioned earlier this model ignores overlapping cases and may over count the contributions of dangles.

### 3.2 “exclusive dangle” model

As mentioned in Section 2.2, accounting for “exclusive dangle” model is non-trivial when dealing with candidates, as ed-candidates do not hold enough information to identify the direction of dangles. To alleviate this problem we provide three different strategies as described below. Each strategy has its pros and cons and should be selected based on the application.

In order to handle the changes for exclusive dangles, we extend the candidate data structure. A candidate base pair, [*i, j*] as implemented in SparseMFEFold holds *i*, the start position, and the energy *V* (*i, j*), as a tuple (*i, V* (*i, j*)) and was stored at the *j*th index of the candidate list. Our extensions to candidate structures involves including the energy values for *W* and *WM* in the candidate tuples as (*i, V* (*i, j*), *W* (*i, j*), *WM* (*i, j*)). The modification reflects the need to store more information about the dangles positions and directions.

### Strategy 1: Trace Arrow implementation

As the first strategy to trace an ed-candidate to its position, we used modified trace arrows. We refer to this strategy as **SparseRNAFolD-Trace**.

Recall that in SparseMFEFold, a trace arrow structures were introduced to identify energy matrix entries that are necessary for calculating energy of internal loops but are not kept as candidates. Here, we define ***ed-trace-arrows*** to hold information about dangle positions to aid with the traceback procedure from ed-candidates. In particular, in the sparse fold reconstruction procedure of SparseRNAFolD, an ed-trace-arrow is checked for a chosen ed-candidate within *W, WM*, and *WM* 2 to adjust the energy and position of the base pair as required. The drawback of this strategy comes from the innate inefficiencies of the trace arrows, meaning increase in space usage. Recall that within SparseMFEFold, we used strategies such as garbage collection and trace arrow avoidance to save space. These strategies are not, however, possible for SparseRNAFolD-Trace, as an ed-candidate cannot be excluded from the optimal MFE path and an ed-trace-arrow is therefore, required for every ed-candidate.

### Bit encoding

Within the second and third strategies as explained next, we employed bit encoding and bit decoding to store the information about the dangle within the energy values to reduce the space usage. Currently, energy values are stored as 32-bits *int* data type. We note that the maximum expected bit usage for energy value of an RNA sequences of up to 20000 base is about 13 bits. We employed a bit shift to store the dangle type in the first two bits of the *V* entries, referred to as *V*_*enc*_, and represented in eq. 8.

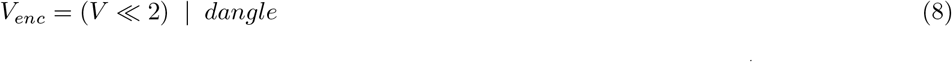

Bit decoding technique was used to retrieve the energy value and type/direction of dangle contributions. Bit decoding was done in two steps. Shifting the encoded energy, *V*_*enc*_, two bits forward gave back the energy, *V* (see eq. 9).

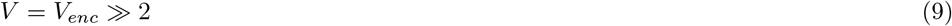

The dangle type is found in the first two bits; no dangle is represented with a “00” in bits, a 5’ dangle with a “01”, a 3’ dangle with a “10”, and a 53’ dangle, with a “11”. The dangle type is decoded using a bit-wise AND with “11” to only keep the first two bits of the encoded energy, as represented in Eq. 10.

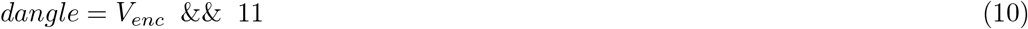

### Strategy 2: Bit encoding with candidate extension

As the second strategy, we used bit encoding within the *W* and *WM* entries of the ed-candidate data structure. We refer to this strategy as **SparseRNAFolD-standard**. This implementation of bit encoding was utilized in *W* and *WM* entries, as other loop types do not deal with dangles.

### Strategy 3: Bit encoding with altered candidate

As the third strategy, we further optimize for space by reducing the candidate size. To reduce candidate size, we stored energy values in ed-candidates in *W* and *WM* as *V* ^d^ minus the dangle energy. We refer to this strategy as **SparseRNAFolD-Triplet**. This strategy allows for correct identification of dangle types regardless of energy parameters used. Note that currently in Turner 2004 energy model, parameter values for 53’ dangle for an external loop and multi-loop are the same. These values may be further estimated and revised in future energy models. The extra calculations to retrieve the *V* ^d^ value ensure accuracy of result in case of such change.

### 3.3 Compared methods

To evaluate performance of our SparseRNAFolD we compared it to two of the best performing methods for prediction of pseudoknot-free RNA secondary structure, namely RNAFold [9] and LinearFold [8].

#### 3.3.1 RNAFold

RNAFold is part of the Vienna RNA package [9]. As discussed in Section 2.1, RNAfold is an *O*(*n*^3^) time and *O*(*n*^2^) space algorithm. It takes an RNA sequence as input and provides the MFE structure as output. RNAFold is well-maintained and highly optimized and is used here as a benchmark for a fast implementation of Zuker and Steigler type MFE algorithm.

#### 3.3.2 LinearFold

LinearFold [8] is a pseudoknot-free RNA secondary structure prediction algorithm which uses heuristic techniques to run in linear time and space. As the main goal of sparsification is to speed up time and space complexity of MFE prediction, we set to investigate how our SparseRNAFolD compares in practice to LinearFold with better asymptotic complexities.

Linearfold employs two techniques to reduce its time and space complexity to *O*(*n*), namely ***beam pruning*** and ***k-best parsing***. Both methods aim to prune the structure path to optimal cases only. Beam pruning works by only keeping a predetermined number (specified by the beam width, *b*) of the optimal states. Within LinearFold, best sets are kept for each possible loop type as defined in the Zuker algorithm: hairpin, multi-loop fragments, and internal loop. Through beam pruning, time complexity is reduced to *O*(*nb*^2^) and the space to (*O*(*nb*) where *b* is the beam width. K-best parsing further reduces the time to *O*(*nblog*(*b*)). We note that due to the heuristic nature of the LinearFold algorithm, it does not guarantee finding the MFE structure for a given RNA sequence.

## 4 Experimental Design

We implemented SparseRNAFolD in C++. All experiments were performed using an Azure Virtual machine. The virtual machine contained 8 vCPUs with 128 GiB of memory.

### 4.1 Dataset

We used the original dataset from SparseMFEFold [37]. This dataset is comprised of 3704 sequences in 6 different families selected from the RNAstrand V2.0 database [1]. The smallest sequence is 8 nucleotides long while the largest is 4381 nucleotides long.

### 4.2 Energy Model

We used the energy parameters of Turner 2004 energy model [21, 32] as implemented in the ViennaRNA package [9].

### 4.3 Accuracy Measures

The number of *true positives* (TP) is defined as the number of correctly predicted base pairings within the structure. The number of *false positives* (FP), similarly, is the number of predicted base pairs that do not exist in the reference structure. Any base missed in the prediction that corresponds to a pairing in the reference structure is a *false negative* (FN).

We evaluate the performance of algorithms based on three measures: sensitivity, positive predictive value (PPV), and their harmonic mean (F-measure).

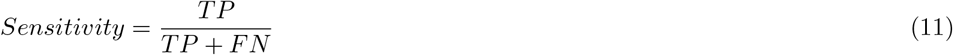

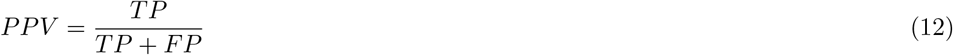

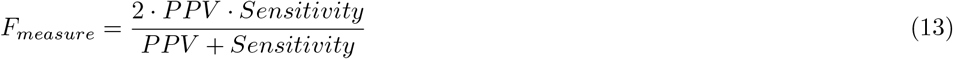

### 4.4 Proof of concept with RNAFold

As the proof of concept for correct implementation of dangle energy models (i.e. “always dangle” and “exclusive dangle”), we assessed SparseRNAFolD against RNAFold. As the MFE structure may not be unique, we restricted our assessment to the MFE value obtained by each method. We found that the MFE predicted by SparseRNAFolD and RNAFold to be the same. Details of the results can be found in our repository.

## 5 Results

Experiments were performed on an Azure Virtual machine. The virtual machine contained 8 vCPUs with 128 GiB of memory. We measured runtime using user time and memory using maximum resident set size.

### 5.1 Alternative Models

We start by comparing the three different strategies in implementation of SparseRNAFolD. SparseRNAFolD-standard was found to be in the middle in terms of memory and time. The effect of additional trace arrows in SparseRNAFolD-Trace had a 27% increase in memory usage on the largest sequence compared to SparseRNAFolD-Standard. However, the increase in computation from the bit encoding only resulted in a 5% increase in time on the largest sequence. We find a similar effect when comparing SparseRNAFolD-standard and SparseRNAFolD-Triplet. The altered triplet structure reduced the memory by 9% but increased the time by 10% due to extra computation. These are highlighted in figure 2.

**Figure 2.**
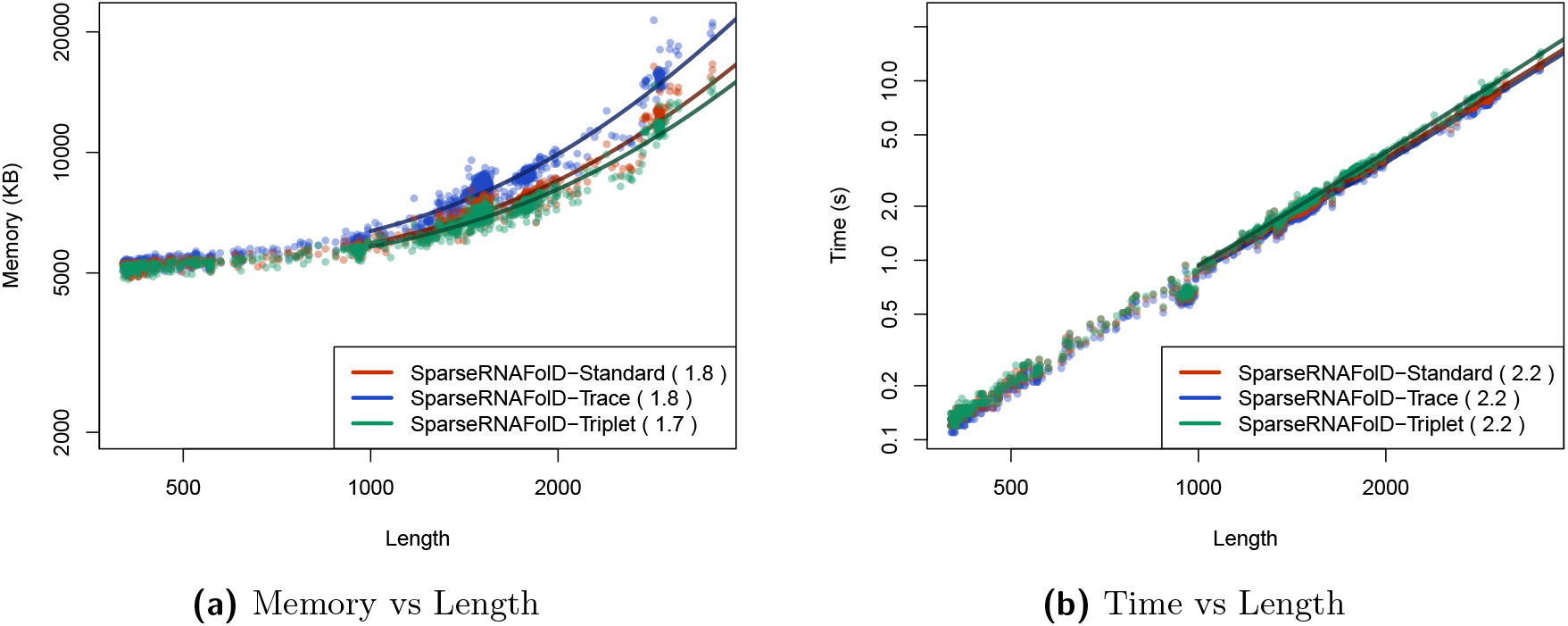
We plot the results of the three versions of SparseRNAFolD when given RNA sequence only as input against and an “exclusive dangle” model each other based on the dataset. (a) Memory Usage (maximum resident set size in KB) versus length (log-log plot) over all benchmark instances. The solid line shows an asymptotic fit (*c*1 + *c*2*n*^*x*^) for sequence length n, constants c1,c2, and exponent x for the fit. We ignored all values < 1000. (b) Run-time (s) versus length (log-log plot) over all benchmark instances. For each tool in both plots, we report (in parenthesis) the exponent x that we estimated from the benchmark results; it describes the observed complexity as Θ(*n*^*x*^).

### 5.2 Comparison with RNAFold

We selected 81 sequences from our dataset with size greater than or equal to 2500. The sequence with the maximum length in the set was 4381 nucleotides long.

As seen in Table 1, while SparseRNAFold’s runtime is comparable to RNAfold, its memory consumption is about five times lower.

**Table 1.** We tabulate the results of the comparison between RNAFold and SparseRNAFolD when given only sequences with length > 2500 from our dataset as input. We looked at time (s) and memory (maximum resident set size in KB) for the minimum, median and maximum length sequence within the constrained dataset.

|  | Run-time (s) |  | Memory: resident set size (KB) |  |
| --- | --- | --- | --- | --- |
|  | RNAfold | SparseRNAFold | RNAfold | SparseRNAFold |
| Minimum | 5.04 | 5.36 | 40148 | 8832 |
| Median | 7.28 | 7.86 | 51284 | 12592 |
| Maximum | 22.08 | 19.94 | 109040 | 16836 |

### 5.3 Comparison with Linearfold

When comparing SparseRNAFolD with LinearFold, we look at the “always dangle” model as LinearFold does not implement the “exclusive dangle” model. We first compared the two algorithms by their predictive accuracy (F-measure).

For comparison, we selected all sequences from our dataset whose structure was available on RNAstrand. We further constrained it to sequences which contained hairpins greater than 3 and no pseudoknots. This resulted in 986 sequences. We found that SparseRNAFolD had a marginally better, but not significant, average F-measure of 0.6394 compared to 0.6391 of LinearFold.

We then assessed their time and space usage. To increase the size of our dataset for this testing, we included a dinucleotide shifted version of our dataset in our test data. We then constrained the size of sequences to those > 400. The max time and memory used by Linearfold on this dataset was 3.34 seconds and 118, 848 KB. In contrast, the maximum time and memory spent by SparseMFEFold was 10.86 seconds and 13, 000 KB. This is illustrated in Figure 3b and 3a. The results show that SparseRNAFolD uses far less memory on even the largest pseudoknot-free sequences in our dataset. Note that the maximum resident set size is nine-times lower than LinearFold. LinearFold, whose time complexity is *O*(*Nb* log(*b*)), did perform faster than SparseRNAFolD as the length of the sequence increased. However, we did find that SparseRNAFolD outperformed LinearFold in practice for sequences of up to about 1000 nucleotides

**Figure 3.**
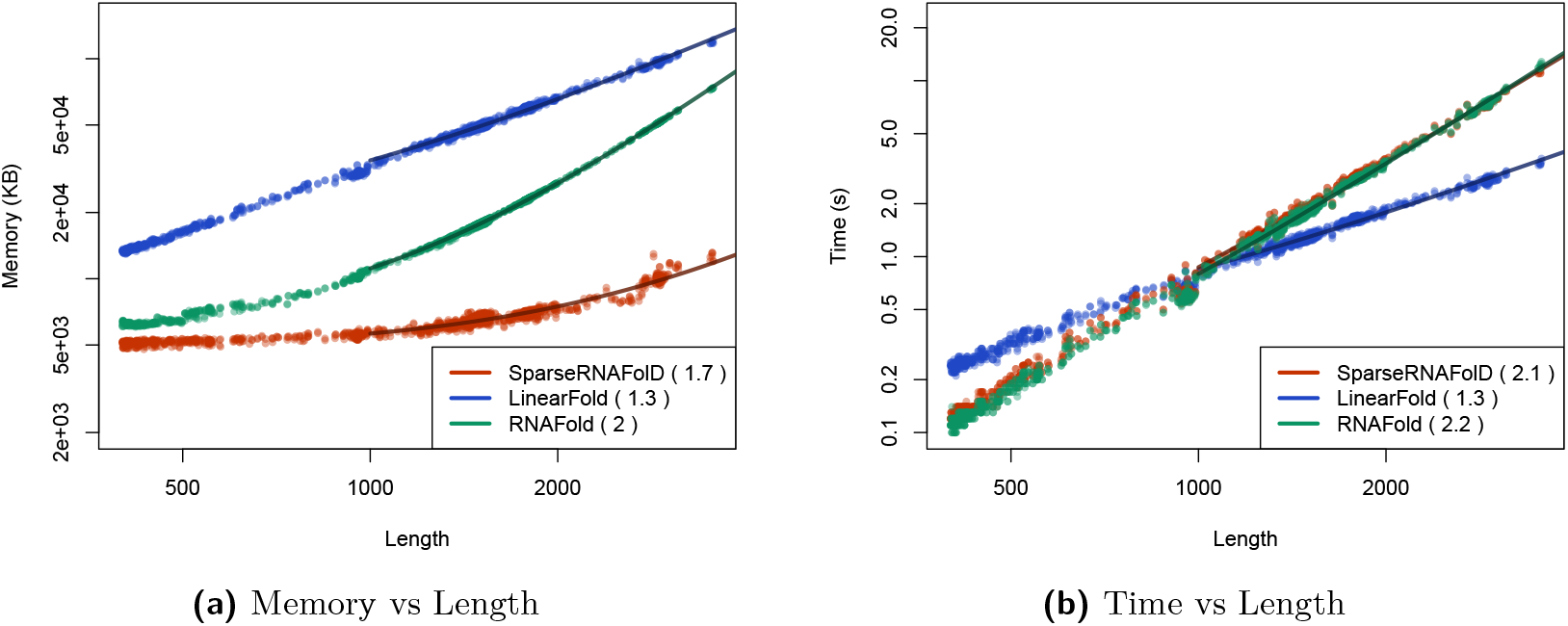
We plot the results of SparseRNAFolD against two state of the art algorithms: RNAFold and LinearFold when given RNA sequence only as input and an “always dangle” model against each other on our dataset and the dinucleotide shuffled version of our dataset. (a) Memory Usage (maximum resident set size in KB) versus length (log-log plot) over all benchmark instances. The solid line shows an asymptotic fit (*c*1 + *c*2*n*^*x*^) for sequence length n, constants c1,c2, and exponent x for the fit. We ignored all values < 1000. (b) Run-time (s) versus length (log-log plot) over all benchmark instances. For each tool in both plots, we report (in parenthesis) the exponent x that we estimated from the benchmark results; it describes the observed complexity as Θ(*n*^*x*^).

### 5.4 Folding with Hard Constraints

As partial information on structure has become more available and is extensively used for better prediction of possibly pseudoknotted structures [13, 10], we further extend our evaluation of the SparseRNAFolD versions to cases where we are folding with a hard constraint [18] in addition to the RNA sequence.

To do so, for each sequence, a pseudoknot-free input structure was generated. The structure was generated by taking two random indices at a time from the sequence. If the two bases could pair, were minimum 3 bases apart, and did not form a pseudoknot with the other base pairs, the base pair was added to the input structure. In order to avoid overpopulating the input structure, the number of base pairs in an input structure was capped at 0.5 × log_2_(*length*). This resulted in an average of 3-7 base pairs per sequence. There was a noticeable decrease in time and space when an input structure was provided in addition of an RNA sequence. Between RNA sequence only as input and sequence as well as an input structure, SparseRNAFolD saw a 67% decrease in time and a 40% decrease in memory. As the input structure reduced the number of candidates for a sequence, the difference in memory was less apparent between the models. SparseRNAFolD-standard had a 6% increase in time from SparseRNAFolD-Trace but a 15% decrease in memory on the largest sequence. From SparseRNAFolD-standard to SparseRNAFolD-Triplet, there was a 8% decrease in memory but a 13% increase in time. Note, even when reducing the number of candidates, the increase in time from Standard to Triplet was greater by 3%. This can be seen in Figure 4.

**Figure 4.**
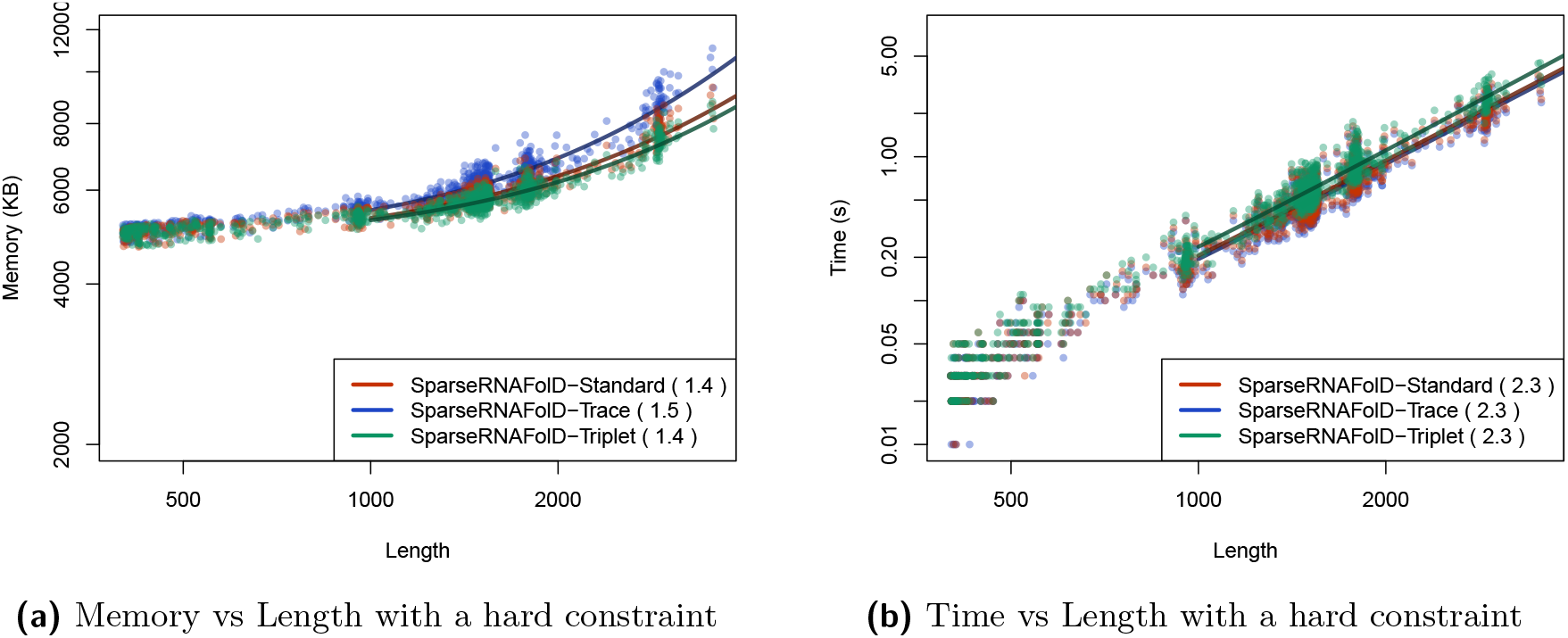
We plot the results of the three versions of SparseRNAFolD when given an RNA sequence, an “exclusive dangle” model, and a random pseudoknot-free structure as input against each other based on our dataset. (a) Memory usage (maximum resident set size in KB) versus length (log-log plot) over all benchmark instances. The solid line shows an asymptotic fit (*c*1 + *c*2*n*^*x*^) for sequence length n, constants c1,c2, and exponent x for the fit. We ignored all values < 1000. (b) Run-time (s) versus length (log-log plot) over all benchmark instances. For each tool in both plots, we report (in parenthesis) the exponent x that we estimated from the benchmark results; it describes the observed complexity as Θ(*n*^*x*^).

## 6 Conclusions

In this work, we introduced SparseRNAFolD, a sparsified MFE RNA secondary prediction algorithm that incorporates dangles contribution to the energy calculation of a sparsified method. We showed that while “no dangle” and “always dangle” models were easy to incorporate to the existing algorithms, “exclusive dangle” introduces non-trivial challenges that need calculated changes to the sparsified recursions to alleviate. We identified three strategies to implement dangle contributions, namely SparseRNAFolD-Trace which utilizes additional trace arrows, SparseRNAFolD-standard, which incorporates bit encoding as well as extension to the definition of candidate structures, and SparseRNAFolD-Triplet, which similar to the SparseRNAFolD-standard utilizes bit encoding, but modifies candidates energy calculation in expectation of possible change of parameters in future. Comparing these three versions on a large dataset, we concluded that SparseRNAFolD-Triplet implementation is the most efficient in terms of memory, and SparseRNAFolD-Trace is the most efficient in terms of time. These two versions showcase how space and time trade-offs can improve the performance based a specific application. The SparseRNAFolD-standard version provides a middle ground for improvement in both time and space, and has been chosen as the standard implementation of our algorithm. While guaranteeing the MFE structure and matching the energy of RNAFold, our SparseRNAFolD is on par with LinearFold on memory usage and run time for sequences up to about 1000 bases. This provides a promising starting to bring dangles contributions to pseudoknotted MFE structure prediction methods in which memory usage is the prohibitive factor [15].

We further assessed the effect of hard structural constraints on performance of SparseR-NAFolD, presenting significant improvement both in terms of time and space. We believe the significant improvement in time and space due to limiting search space by hard structural constraints can have a more pronounced impact on sparsified pseudoknotted MFE prediction, which is our ultimate goal.

Finally, memory consumption becomes a bottleneck for prediction of MFE structure for long RNA sequences or MFE pseudoknotted structure prediction. Utilizing power of computational servers such restrictions have been somewhat alleviated. Sparsification provides improvement in both time and space requirements and can be used to bring back computations to personal computers providing equal access to the existing technology. In addition, improvement in memory usage, can improve use cases of computing cluster as assigned memory to a computing node is also limited.

## 7 Author contributions statement

MG and HJ conceived the experiment(s), MG. conducted the experiment(s), M.G. analysed the results. M.G. and H.J. and S.W. wrote and reviewed the manuscript.

